# Effects of overt and covert attention on decision-making dynamics in prefrontal cortex

**DOI:** 10.64898/2026.05.18.723036

**Authors:** Nathan T. Munet, Joni D. Wallis

## Abstract

Value-based decision-making is a dynamic and idiosyncratic process which requires appraising the value of options and comparing the values to select the best choice. An emerging view of orbitofrontal cortex (OFC) is that it achieves this by representing values serially during deliberation, alternating back-and-forth between transient states that encode the value of different options. While the time spent in each value state is known to reflect choice behavior, the source of the alternating dynamics remains unclear. One possibility is that fluctuations in value may be driven by attentional shifts between the choice options. Conversely, value dynamics in OFC may be generated locally, enabling OFC value signals to influence decision-making independently from attention. To test these hypotheses, we recorded from OFC and lateral prefrontal cortex (LPFC), a major attentional area in prefrontal cortex, to determine whether their population-level activity correlated in a manner consistent with crosstalk between neuronal systems involved in value and spatial attention. We found that OFC and LPFC selectively encoded option values and spatial locations, respectively, reflecting their specialized roles in cognition. Despite this functional dissociation, both OFC and LPFC dynamics were strongly affected by overt attention: which caused the value and spatial location of the fixated option to be represented at the same time. Additionally, fluctuations in the encoding strength of value in OFC and space in LPFC were temporally correlated above and beyond the effect of gaze, reflecting the effect of covert attention.

## Introduction

During value-based decisions, orbitofrontal cortex (OFC) represents available option values discretely and dynamically, alternating between value-encoding states pertaining to each of the options at hand ^1^. While these ‘flip-flopping’ value states do not directly signal the identity or the timing of the chosen response, they are thought to influence choice via integration by downstream regions involved in action selection. The existence of an underlying integration mechanism is supported by the observation that both the frequency and the amount of time OFC spends in each option’s state predict the speed and outcome of the decision ^1^. Furthermore, value coding in accelerates the rate of response-encoding ramping activity in anterior cingulate cortex ^2^, suggesting that anterior cingulate cortex could be an important target for OFC outputs in driving choice behavior

While these findings shed light on the functional output of OFC’s value signal, less is known about the source of this signal’s flip-flopping dynamics. Specifically, it remains unclear whether value flip-flopping arises from local recurrent dynamics within the value network, analogous to the mechanisms underlying perceptual multistability ^3^, or whether it reflects global shifts in selective attention between choice alternatives. Toward the latter hypothesis, the flip-flopping motif in OFC bears some resemblance to alternating state dynamics reported in a variety of visual and prefrontal areas involved in attention and working memory ^4–6^. Thus, it is feasible that the dynamics of value coding in OFC could be the signature of more general attentional dynamics coordinated across a variety of brain regions processing different task information in parallel ^7^. While early examination of single-trial value dynamics in OFC found no relationship between overt attentional shifts and value flip-flops ^1^, more recent studies show that OFC’s value code can be influenced by gaze ^8–11^.

In this study, we aimed to quantify how both overt eye movements and covert visuospatial attention might affect value dynamics in OFC. To assess the role of gaze in OFC’s value representations, we asked whether value information decoded from OFC was consistent with subjects’ gaze location and whether shifts in the value code were linked to shifts in gaze. We additionally measured the dynamics of covert attention by recording from lateral prefrontal cortex (LPFC) simultaneously with OFC, since LPFC is a critical area in orchestrating top-down attention across the brain ^12,13^, and the dynamics of the ‘spotlight’ of attention ^14^ can be decoded from LPFC on single trials ^15,16^. While LPFC is not directly connected to OFC ^17^, it does reliably track network-level attentional signals, allowing us to utilize the spatial signals in LPFC as a proxy for covert attention and test whether the dynamics of covert attention affect the dynamics of value in OFC.

## Results

### Value-based choice behavior

To examine whether visuospatial attention affects valuation during choice, we first trained subjects to associate pictures with different reward outcomes (amount and probability of juice reward). We then had our subjects choose between these pictures, based on their reward values, as they were presented at opposite locations on a screen (Figure 1A; locations varied trial-by-trial). Half of these choices were free choices, offering two pictures with distinct reward associations and thus different values. These free choices served two purposes. First, they allowed us to fit subjective value models that quantified how the amount and the probability of reward contributed to choice behavior. Second, they provided a test condition to characterize the dynamics of two-item attentional and evaluative processes in the prefrontal cortex (PFC) and investigate the relationship between these dynamics. The remaining half of choices were forced choices, offering only a single picture option at a single location. The forced choice trials provided a ‘ground truth’ condition to estimate a neural code for value by establishing a relationship between neural activity and the isolated picture values.

**Figure 1.**
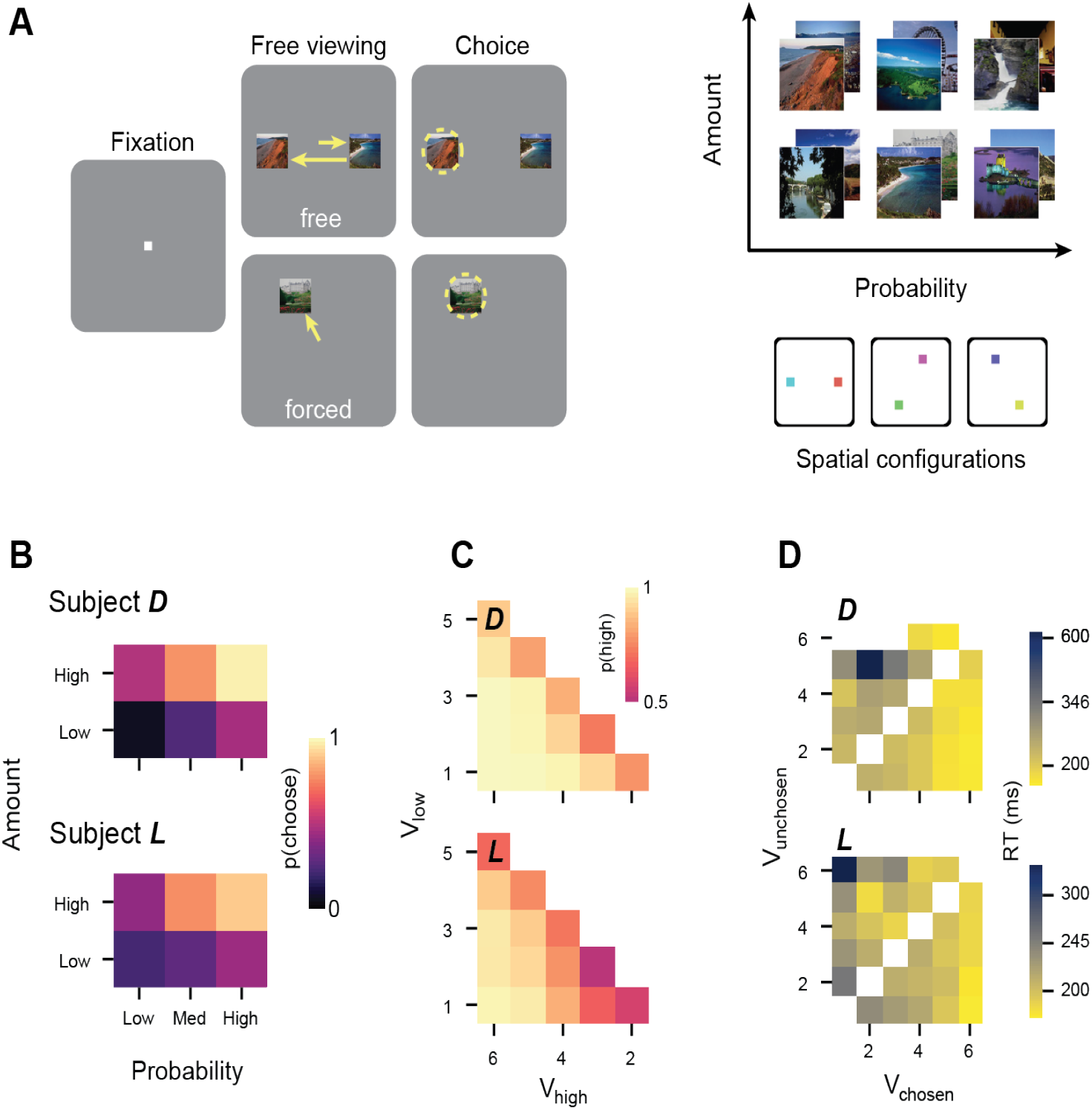
Value-based choice behavior. A) Value-based choice task and reward contingencies. On each trial, subjects were offered either one (forced choice) or two (free choice) pictures which were associated with a specific amount and probability of reward (top right panel). Choices were made by fixating on a picture for 350 ms. B) Choice likelihood, *p*_*choose*_, as a function of reward probability and juice amount. Choice likelihood monotonically increased with both reward variables, indicating that both the probability and amount of reward contributed to choice behavior. C)The likelihood of choosing the higher value option, *p*_*high*_, as a function of the ordinal rank of the higher value (*V*_*high*_) and lower value (*V*_*low*_) option. Both offer values contributed to choice since choice likelihood monotonically increased with *V*_*high*_ and decreased with *V*_*low*_. D) Choice response times (RTs) as a function of the chosen and unchosen values (*V*_*chosen*_ and *V*_*unchosen*_). The color axis is logarithmic. Response times were faster when *V*_*chosen*_ was high and slower when the value difference was small. White squares indicate unsampled data points.

Choice patterns confirmed that both the amount and the probability of reward guided decision-making, indicating that both variables were integrated into value (Figure 1B). To quantify the value function, we fit choice behavior on each session to three different subjective value models, testing whether amount and probability were integrated multiplicatively (as expected value), linearly (with each variable contributing independently to choice as a weighted sum), or as a hybrid of the two, with both linear and multiplicative components. We compared the performance of the models using the Bayesian Information Criterion (BIC). The average BIC was lowest for the hybrid model for both Subject D (Hybrid: 228; Linear: 238; EV: 355) and Subject L (Hybrid: 472; Linear: 496; EV: 548), indicating that the hybrid model yielded the best behavioral fit. We therefore used the value estimates from the hybrid model for all subsequent analyses.

We next assessed the relationship between the offer values and choice behavior. On free choices, subjects reliably chose whichever option was highest in value (Subject D: 92% of trials; Subject L: 80% of trials), and this preference for high-value options was consistent across all choice offerings (Figure 1C). Both the discriminability of the offer values (i.e., how far apart they were in value) and the value level of the higher-value option predicted the likelihood that the higher value would be chosen (stepwise logistic regression; *p* < 0.001 for both regressors in both subjects). Similar trends were reflected in how quickly subjects arrived at a final selection. Strong discriminability between options and more rewarding best offers led to faster choices (Figure 1D).

### Attention behavior

To measure visuospatial attention in PFC during two-item decisions, we interleaved the value-based choice task with a separate attention-guided change detection task in alternating blocks. Similar to how forced choice value trials were a ground truth condition for estimating a value code, the attention trials served as a ground truth for identifying a spatial attention code by allowing us to manipulate where subjects oriented their attention.

Monkeys covertly attended two Gabor targets, presented in the same spatial locations used in the free choice value trials, and performed a saccade towards a target to report which (if any) rotated after a 500-800 ms random delay (Figure 2A). Targets were color-coded to indicate that the green (valid) target was more likely to rotate than the red (invalid) one. This encouraged subjects to preferentially attend to the location of the valid target. To ensure subjects were incentivized to carefully attend to the screen and preclude a default strategy of choosing the valid target regardless of whether it rotated, 25% of attention trials were ‘catch trials’ in which neither target rotated and subjects had to maintain central fixation to earn a reward.

**Figure 2.**
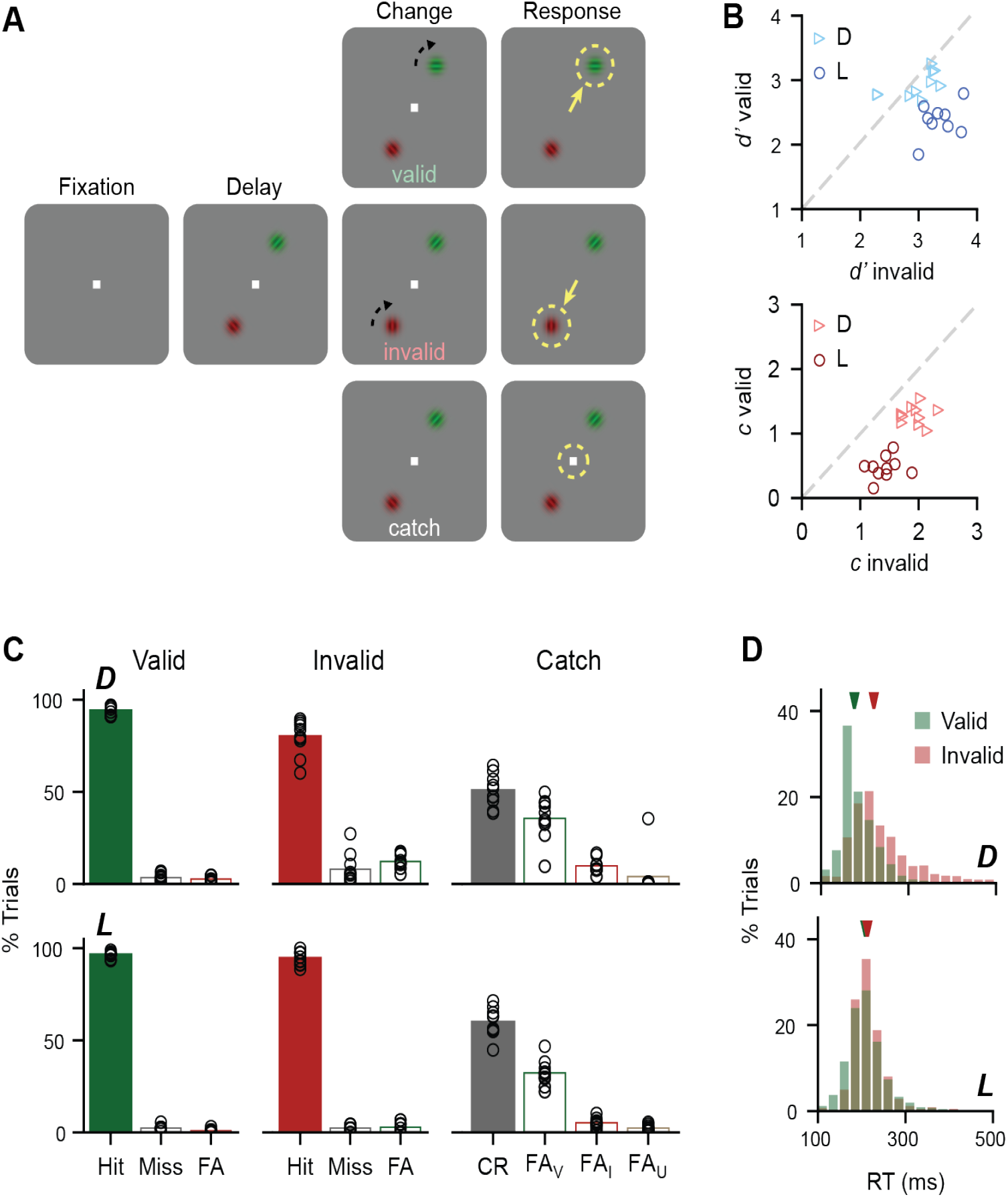
Attention task behavior. A) Subjects were presented with two Gabor grating targets to covertly attend over a variable delay (500 - 800 ms). Subjects reported which target changed (± 25° rotation) after a brief stimulus blink by saccading to the rotated target within 500 ms. They reported ‘catches’ (no rotation; 25% of trials) by maintaining central fixation over the response period. Rotations were more frequent at the green (‘valid’) location than the red (‘invalid’) location. B) Top: Sensitivity, *d’*, toward detecting valid versus invalid changes. Bottom: choice criteria, *c*, for reporting valid versus invalid changes. C) Change detection response rates on valid change (left), invalid change (center), and catch trials (right). Data points indicate single-session response rates (n = 10 sessions for both subjects). Bars indicate means across sessions. Bars are colored according to which target was chosen: green = valid; red = invalid; grey = neither target. Filled-in bars indicate the correct response for each condition. FA = False Alarm; FA_V_ = FA to valid target; FA_I_ = FA to invalid target; FA_U_ = FA to unclassified location. D) Reaction time (RT) distributions for valid (green) and invalid (red) hit trials. Arrowheads indicate median RTs (Subject D: 176 ms valid vs 221 ms invalid hit RT, *p* < 0.001; Subject L: 208 ms valid vs 213 ms invalid hit RT, *p* < 0.001; median RTs, rank-sum test).

We fit a multi-variate signal detection theory (SDT) model to the response rates for each possible outcome (valid change, invalid change, or catch). This model assessed whether response behavior reflected an attentional bias that could be captured by target-specific differences in either the sensitivity to target rotations or the criterion for selecting a given target. This allowed us to quantify the threshold of evidence needed to report a detection. As expected, change detection behavior was biased according to the attentional cues, which the SDT models largely attributed to a significantly decreased choice criterion for the valid target compared to the invalid one (Figure 2B; D: *c*_*val*_ = 1.3 ± 0.1, *c*_*inv*_ = 2.0 ± 0.1, *p* = 0.002; L: *c*_*val*_ = 0.5 ± 0.1, *c*_*inv*_ = 1.4 ± 0.1, *p* = 0.002; rank-sum test). This manifested in biased false alarm (FA) reporting of the valid target over the invalid one (Figure 2C.; D: 35.6 ± 3.6% valid vs. 9.8 ± 1.2% invalid FAs; L: 32.4 ± 2.2% valid vs. 5.3 ± 0.8% invalid FAs), indicating that subjects were primed to saccade to the valid location. Consistent with this priming, we found that reaction times (RTs) were significantly faster when changes occurred at the valid location compared to the invalid location (Figure 2D).

In contrast to the attentional effects on choice criteria, biases relating to perceptual sensitivity were less clear. Per the SDT model, Subject D was similarly sensitive to either target rotating (Figure 2B; *d’*_*val*_ = 2.9 ± 0.1, *d’*_*inv*_ = 3.0 ± 0.1, *p* = 0.49), while Subject L was surprisingly more sensitive to the invalid target (*d’*_*val*_ = 2.4 ± 0.1, *d’*_*inv*_ = 3.9 ± 0.6, *p* = 0.002). Notably, despite this lack of an attention-congruent sensitivity bias, both subjects were highly accurate at reporting valid changes (Figure 2C): Subject D more accurately reported valid changes compared to invalid ones (93.8 ± 0.8% valid hits vs. 79.8 ± 3.0% invalid hits), whereas Subject L performed near ceiling in both conditions (96.5 ± 0.5% valid hits vs. 94.5 ± 1.2% invalid hits). Together, our results indicate that the task successfully modulated attention and did so by biasing response behavior rather than perceptual sensitivity.

### OFC and LPFC are specialized toward encoding either value or space

We performed high-density, simultaneous recordings in OFC (D: N = 1942 neurons recorded in 10 sessions, L: N = 2378 neurons recorded in 10 sessions) and LPFC (D: N = 2238 neurons, L: N = 2444 neurons) using primate Neuropixels electrodes (Figure S1). To determine how neurons in each prefrontal region encoded the values and spatial locations of the pictures presented, we first analyzed single-neuron tuning characteristics on the forced choice trials of the value task. Because forced choices always entailed single pictures offered in isolation, we were able to construct simple regression models to quantify how individual neurons encoded each possible combination of value and location (Figure 3A, B; Figure S2). The regression coefficients revealed that while neurons sensitive to space and value could be found in both regions, OFC neurons preferentially encoded picture value (D: 22% value vs. 4% space, *χ*^*2*^ = 256, *p* < 0.001; L: 20% value vs. 8% space, *χ*^*2*^ = 141, *p* < 0.001), whereas LPFC neurons had a more dominant spatial code (D: 29% space vs. 10% value, *χ*^*2*^ = 271, *p* < 0.001; L: 32% space vs. 17% value, *χ*^*2*^ = 142, *p* < 0.001).

**Figure 3.**
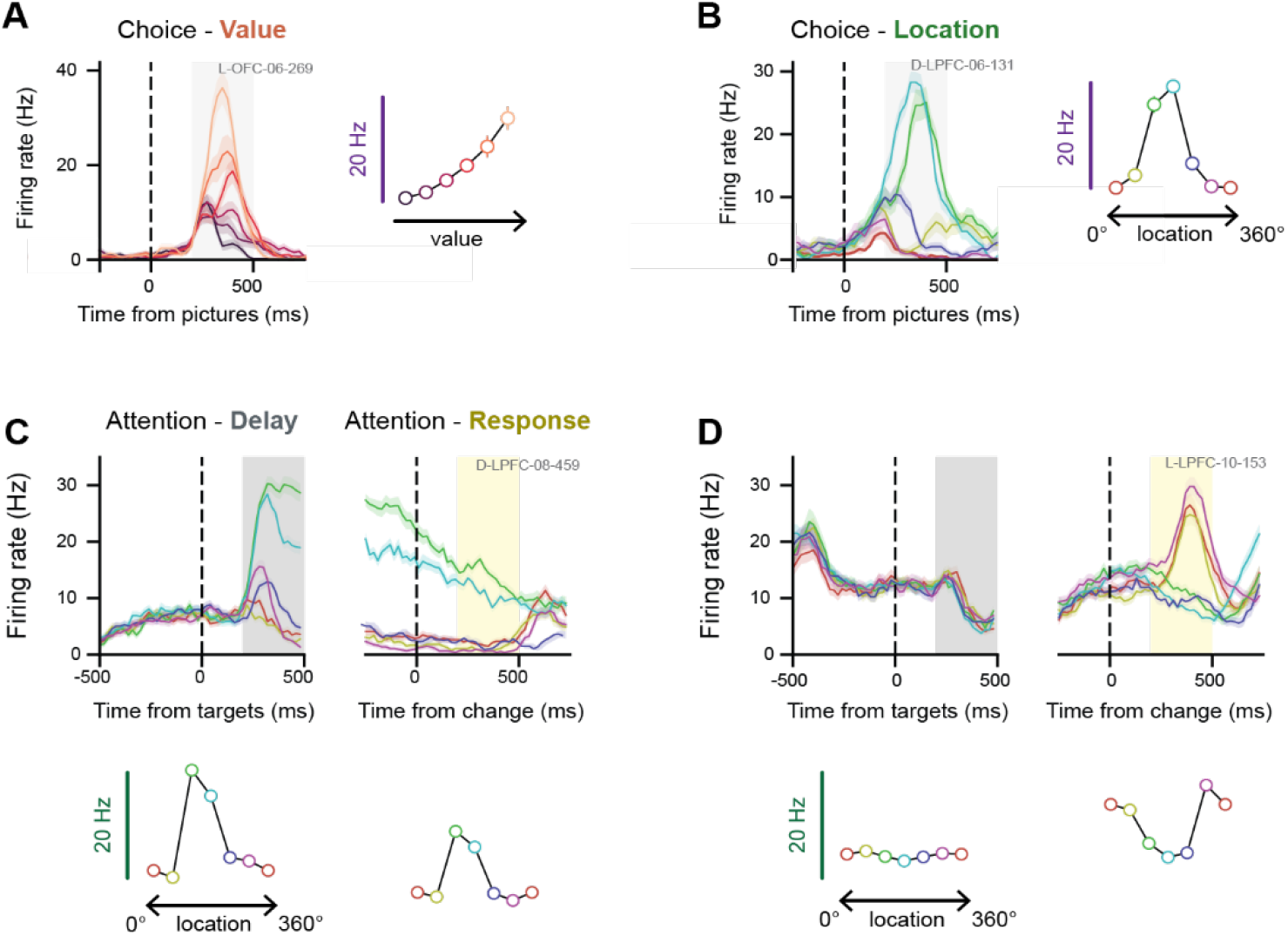
Single-neuron encoding of value and space. A) Spike density histogram of an example OFC neuron encoding value (left) and the corresponding value tuning function (right) during the value-based choice task. B) Spike density histogram of an example LPFC neuron encoding spatial location (left) and the corresponding spatial tuning function (right) during value-based choice. C) Spike density histogram (top) showing an example LPFC neuron that has consistent spatial tuning at the time of target presentation and stimulus change during the attention task. Tuning curves (bottom) were calculated from 200 - 500 ms after target onset and 200 - 500 ms from the stimulus change. D) Same as (C) except showing an example LPFC neuron with spatial tuning only at the time of the change.

The attention task allowed us to better characterize the spatially tuned LPFC neurons. Over the duration of the attention task, LPFC neurons showed heterogeneous spatial tuning profiles (Figure 3C, D; Figure S3). Across the population, the proportion of neurons tuned to the valid target location began to ramp during the delay period and peaked during saccade initiation (Figure S4). In contrast, OFC contained few spatially selective neurons during any epoch of the task.

We next turned to population-level analysis to determine whether dynamics in spatial and value coding coordinated during value-based decisions. To address this question, we trained linear discriminant analysis (LDA) classifiers to decode the picture values and spatial locations from population spiking activity in either OFC or LPFC. The value decoders were trained on firing rate vectors during the forced choice trials of the value task, whereas the spatial decoders were trained on firing rates during the delay period of the attention task. Thus, the two types of decoders were trained on independent task conditions designed to isolate neuronal representations of either picture value or spatial attention. The decoders were then used to estimate distinct value and spatial codes on the free choice trials as subjects freely viewed and deliberated between the two pictures offered.

Our first aim was to determine which task information—values or spatial locations—could be decoded from each region of PFC. We could decode the value of both the chosen and unchosen picture above chance from both areas, but decoding accuracy was significantly higher in OFC than in LPFC (Figure 4A, B). In contrast, the chosen location was readily decodable from LPFC, but there was little information about the picture locations in OFC (Figure 4C, D). Decoding at a finer temporal resolution (80 ms sliding windows) additionally revealed transient information about the unchosen location immediately after picture onset (Figure 4C), although this was not reliably decoded above chance over the coarser 100 - 500 ms analysis window (Figure 4D). Together these initial decoding results point to dissociable functional specializations of OFC and LPFC in decision-making, wherein OFC operates in a value-centric reference frame and LPFC operates in a visuospatial one.

**Figure 4.**
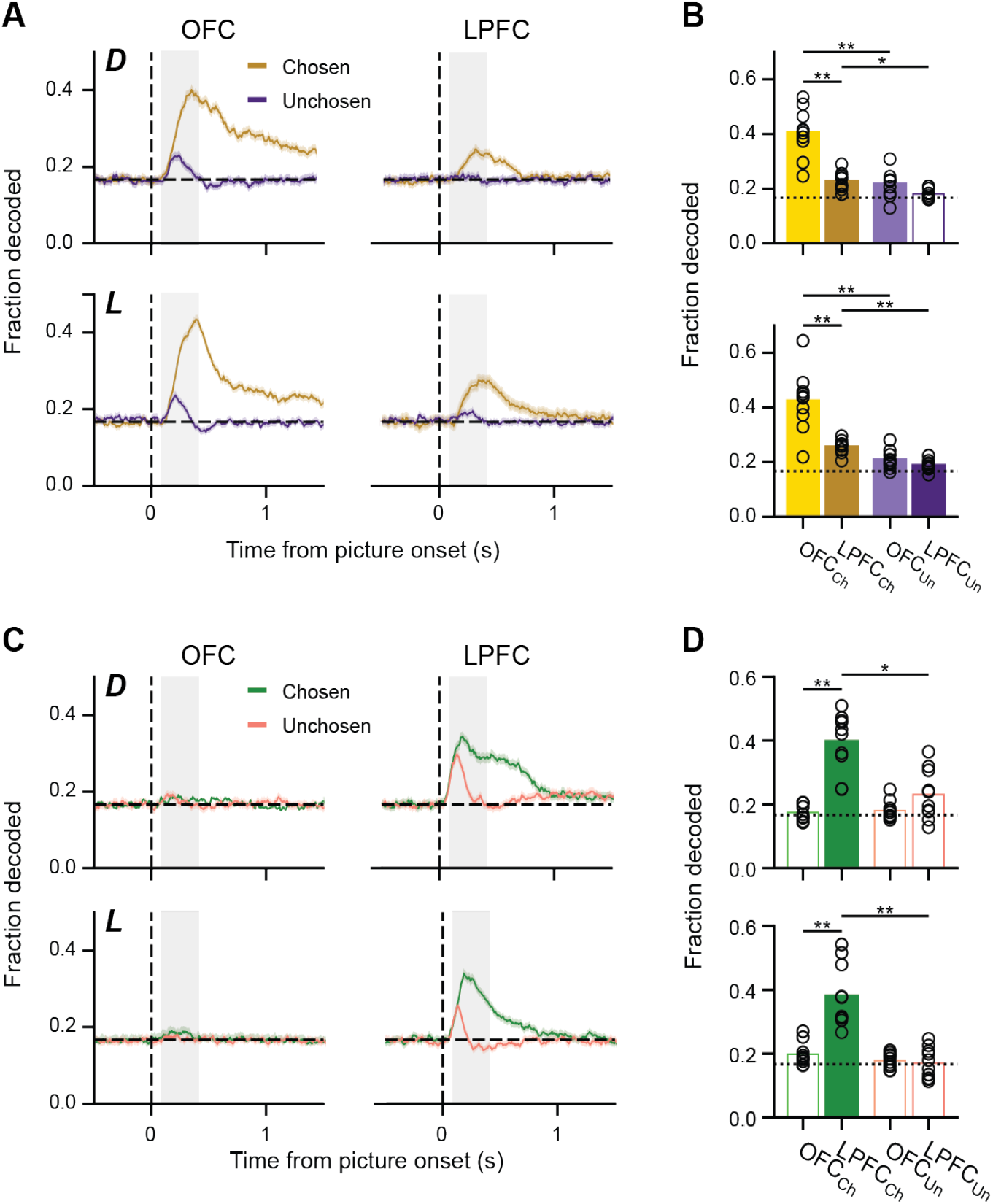
Dissociable population codes for value and space. A) Proportion of trials where we decoded chosen or unchosen values from either OFC or LPFC as a function of time from picture onset (20 ms sliding Gaussian kernel, stepped by 5 ms). Error bounds indicate bootstrapped 95% confidence interval. The grey shading indicates the time range used in (B). B) Proportion of trials where we decoded chosen or unchosen values during the 100 - 400 ms time following picture onset. Data points indicate individual sessions. Values were more frequently decoded from OFC than LPFC, and chosen values were decoded more frequently than unchosen values in both regions. Filled bars indicate that the proportion was significantly greater than chance (one-sided *t*-test, *p* < 0.01 with Bonferroni correction). Asterisks indicate significant differences between groups (paired *t*-test with Bonferroni correction, * *p* < 0.01, ** *p* < 0.001). C, D) Same as (A, B) but showing decoding of chosen or unchosen spatial locations.

### Overt and covert attention affect spatial and value dynamics in PFC

The primary goal of our study was to determine whether value state dynamics in OFC are related to shifts in attention. Prior work from our group was unable to identify a relationship between value states and eye movements ^1^. However, in this earlier study we were limited by the number of neurons from which we could record, and as a result, value decoding was relatively noisy. More recent work has demonstrated clear evidence for gaze-contingent value processing in OFC ^8^. To reconcile these findings and determine the scope of any possible connections between OFC value coding and attention, we assessed gaze dynamics as an indicator of overt attention and asked whether gaze sufficiently explained which option’s value was represented in OFC.

We began by analyzing eye movements during free choice trials. We defined deliberative trials as those in which subjects made more than a single fixation to indicate their choice. Subject D was more likely to engage in overt deliberation than Subject L, making two or more fixations on 29% of choices compared to only a 9% multiple fixation rate for Subject L (Figure 5A). For both subjects, deliberation was significantly more likely when there was a larger difference in value between the alternative option and the currently fixated option (Figure 5B; Table 1). We also noticed that Subject L’s median RT (176 ms) for the initial saccade was slower than that of Subject D (154 ms) which might be consistent with Subject L prioritizing the accuracy of the first saccade such that multiple fixations were not required to make a choice. This also suggests that Subject L may rely on covert attention more than Subject D.

**Figure 5.**
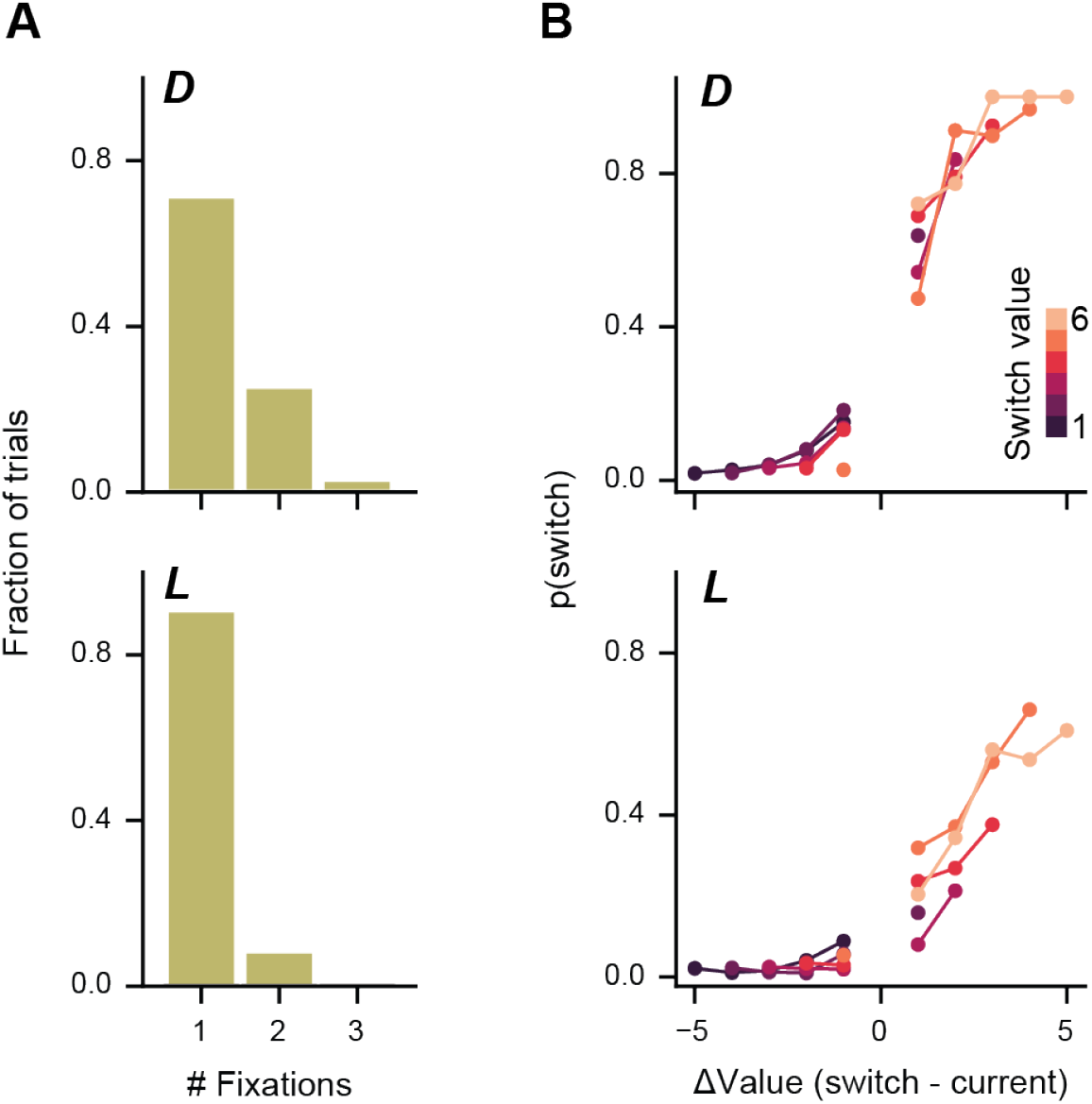
Gaze behavior during choice. A) Number of fixations per trial during free choice trials. B) Probability of switching gaze from the currently fixated picture to the alternative option. Switch probabilities were assessed as a function of the difference in value between the prospective switch target and the currently fixated one (horizontal axis), as well as the value level of the switch target (colored curves, mapped to the ‘switch value’ color bar).

**Table 1.** Effect of option values on deliberation. To determine how the value of the options affected the likelihood of deliberation (defined as fixating more than one option during choice) we performed a logistic regression with parameters of the value difference between the currently fixated option and the alternative option (*β*_*delta*_) and the value of the alternative option to which the eye switches (*β*_*switch*_). For both subjects, the main determinant of deliberation was whether there was a large difference in value between the alternative option and the currently fixated option (*β*_*delta*_).

|  | Subject D |  | Subject L |  |
| --- | --- | --- | --- | --- |
| pseudo $R^2$ | 0.51 | $p \approx 0$ | 0.23 | $p = 1.0 \times 10^{-175}$ |
| $\beta_{\text{delta}}$ | 2.8 | $p = 1.8 \times 10^{-241}$ | 1.3 | $p = 3.6 \times 10^{-78}$ |
| $\beta_{\text{switch value}}$ | -0.33 | $p = 6.5 \times 10^{-9}$ | 0.17 | $p = 6.6 \times 10^{-3}$ |
| constant | -1.7 | $p = 5.1 \times 10^{-186}$ | -3.0 | $p \approx 0$ |

We next examined how OFC value representations varied over this behavior using our value decoder (Figure 6A). On single fixation trials, representation of the fixated value dominated over that of the unfixated value, and the likelihood of decoding the unfixated value fell to chance (defined as the same likelihood as decoding an unavailable option) roughly 160 ms after fixation onset. On deliberative trials, when the subject switched their gaze, the value code updated to reflect the newly fixated picture (Figure 6B). We performed the same analysis on the spatial representations in LPFC (Figure 6C). On single fixation trials, LPFC’s spatial code was similar to OFC’s value code in that the current fixated location dominated over the unfixated one around the time of fixation. However, unlike value, which lagged fixations, spatial information emerged before fixation onset, consistent with LPFC’s role in guiding saccades and attentional shifts. On multi-saccade trials, the two subjects showed somewhat different spatial dynamics with respect to gaze (Figure 6D). For Subject D, the spatial decoder tracked the location of the currently viewed option, similar to the value dynamics. In contrast, for Subject L, the location of the higher value option was persistently dominant over that of the lower value option, regardless of which option was initially fixated.

**Figure 6.**
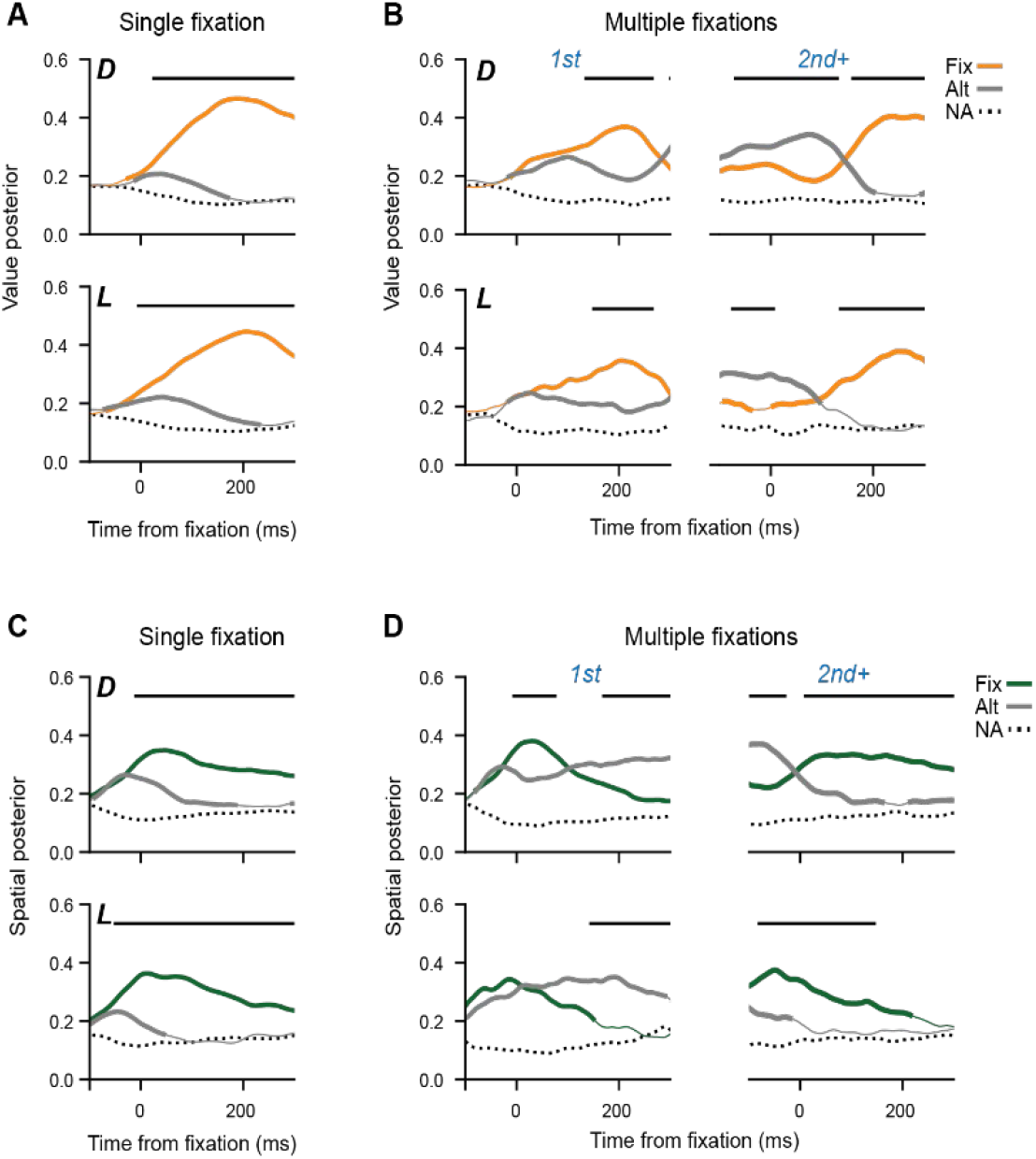
Gaze-contingent dynamics in value and spatial codes. A) OFC value decoder posterior probabilities (mean ± SEM) for fixated, alternative, or unavailable options during trials with single fixations or multiple fixations. Thick lines indicate a significant difference between the fixated or alternative option as compared to the unavailable option. Horizontal bars indicate significant differences between the fixated and alternative option. Significance was assessed using paired *t*-tests with Bonferroni correction evaluated at *p* < 0.01. B) Same as (A) but showing LPFC spatial decoder posterior probabilities.

Although we did not see fixation-incongruent value representations when we averaged across trials, they could potentially be more apparent on individual trials. To examine this possibility, we decoded value in a fixed window 160 - 300 ms after fixation onset, which was when the value representation typically corresponded to the value of the fixated option. In Subject L, we were frequently able to decode the unfixated option in addition to the fixated option (Figure 7), but this was not the case for Subject D. This supports our conclusion from the eye movement behavior (Figure 5) that Subject D relied more on overt eye movements during decision-making, whereas Subject L relied more on covert attention.

**Figure 7.**
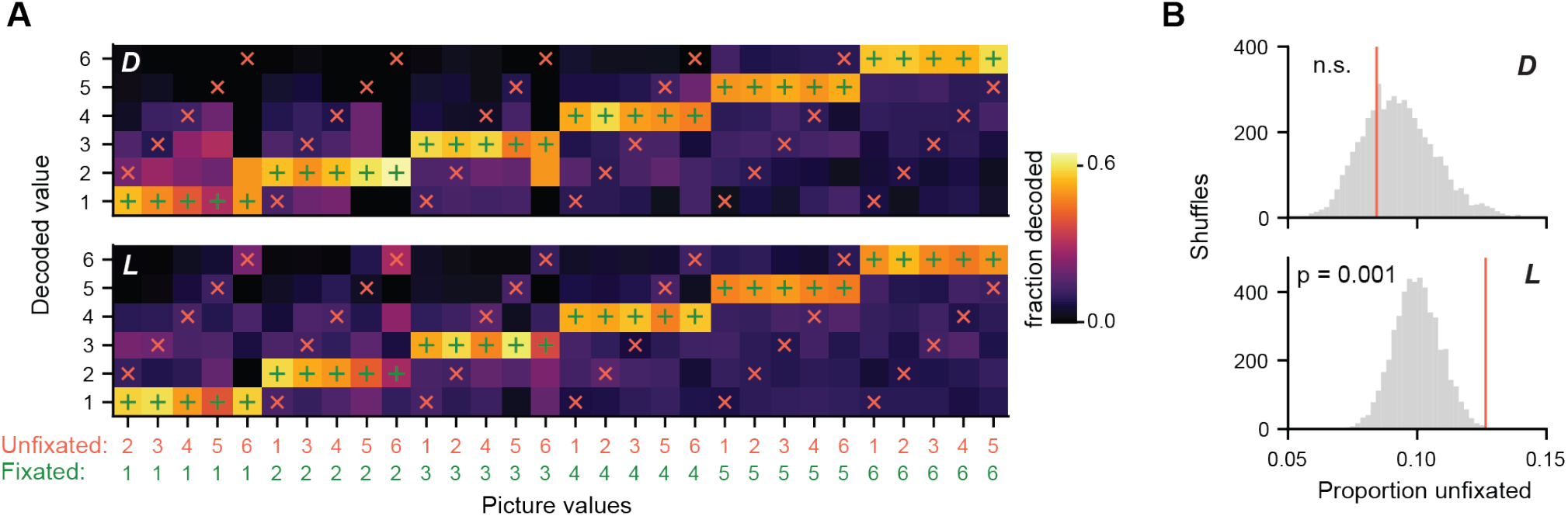
Value representations of fixated and unfixated choice options. A) Confusion matrix showing the fraction of fixations for which each possible value was decoded for each possible combination of fixated and unfixated options during the late phase of each fixation (200 - 300 ms after fixation onset). B) Red vertical line indicates the fraction of fixations for which the unfixated value was decoded (red line). Significance was assessed against a null hypothesis that the probability of decoding an unfixated value comes from the same distribution as that of decoding one of the values that was not available on a given trial (grey histogram; 5000 permutations per subject). Unfixated values were significantly more decodable than unavailable ones for Subject L (*p* = 0.001) but not for Subject D, showing that, at least for one of our subjects, the value code reflected information that could not be explained exclusively by gaze.

While gaze explained much of the variability in OFC’s value representations, we were interested in whether covert attention could play an additional role in guiding value dynamics. To address this question, we asked whether OFC value states and LPFC spatial states were correlated. When we cross-correlated the posteriors from the value and spatial decoders after removing the trial-averaged gaze dynamics, we found little evidence of a relationship between the representations of the fixated location and the fixated value, but we did find stronger evidence for a positive correlation between the spatial and value codes for the unfixated option. These findings suggest that there may be coordinated covert sampling of the unfixated option’s value and location in PFC (Figure 8). The cross-correlations were significant over a larger portion of the fixation window on single fixations (Figure 8A) compared to ‘switches’ (i.e., follow-up fixations when multiple fixations were made; Figure 8B), potentially indicating more robust coordination in the absence of overt deliberation. This result is consistent with the idea that covert attentional states may drive value dynamics in lieu of overt attention.

**Figure 8.**
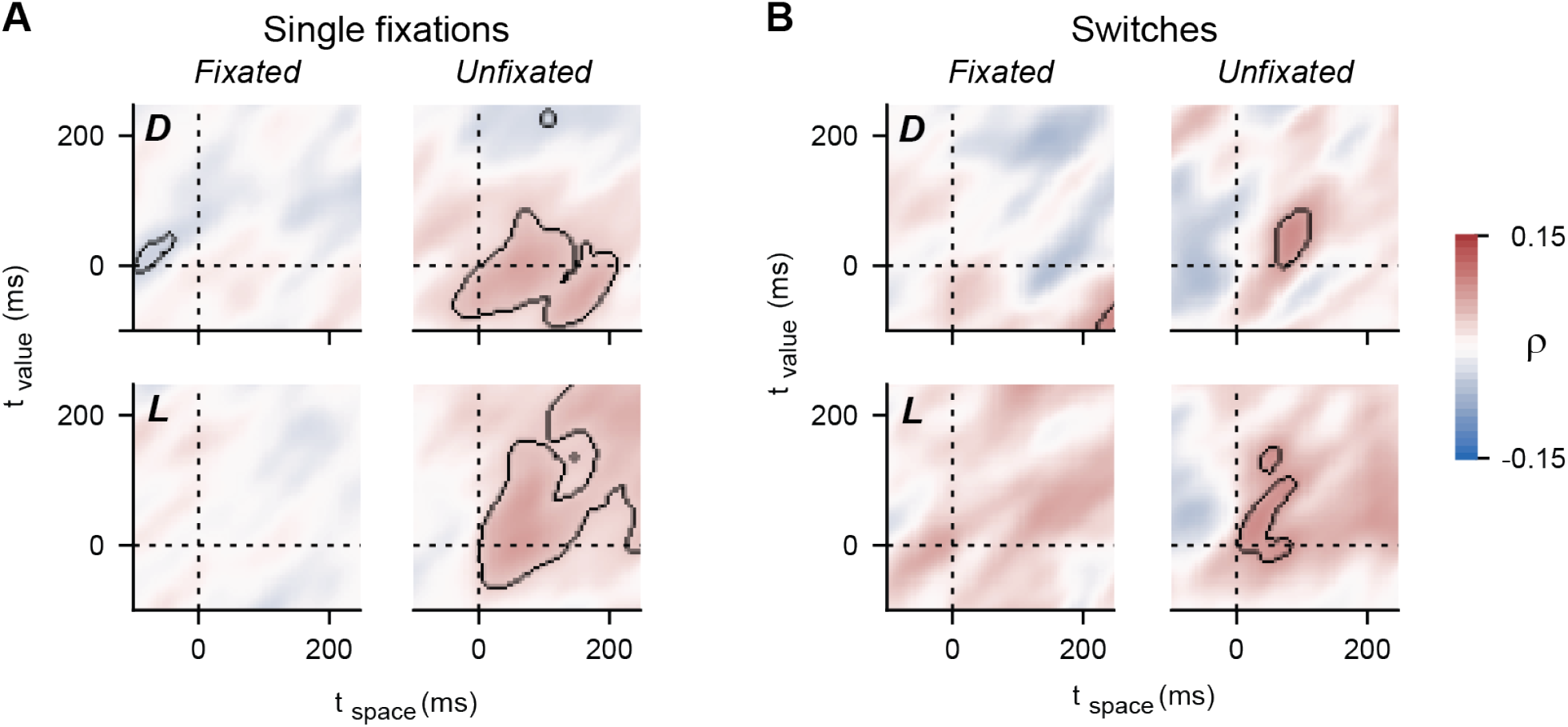
Cross-correlation between LPFC spatial and OFC value representations. A) Cross-correlation, *ρ*, between the posteriors from the LPFC spatial decoder and the OFC value decoder posteriors for the fixated or unfixated picture on single-fixation trials. Decoder time series were aligned to fixation onset and cross-correlated (100 ms time window with 5 ms step size). Black lines denote significant cross-correlations (*p* < 0.01, permutation test). B) Same as (A) but aligned to fixational switches (i.e., fixations where gaze was switched from one picture to the other).

Lastly, we considered the possibility that saccades toward the pictures might be rhythmically entrained to oscillations in the local field potential (LFP), which could mediate coordinated coding across PFC. To examine this, we used linear regression to predict the time of first fixation based on the time of the first theta, alpha, or beta peak following stimulus onset. We were unable to identify any consistent relationships between the fixation timing and LFP oscillations (*p* > 0.05 for both subjects in both regions for each LFP band, except for Subject L in OFC’s alpha band which was weakly significant at *p* = 0.02 and *r*^*2*^ = 0.0004).

## Discussion

We found evidence of coordination between two anatomically and functionally dissociable subsystems within the prefrontal cortex during a value-based decision-making task. At both the single-neuron and population level, OFC encoded value, whereas LPFC encoded visuospatial information. Despite this segregation, processing within the two areas is tightly coordinated during decision-making via both overt eye movements (i.e., gaze) and covert attention. This ensures that the two areas represent the same choice option at the same time as individual decisions are made, allowing goal-related processing to interface with the decision-maker’s perception of and interactions with the external environment.

It has long been understood that LPFC neurons have strong spatial selectivity subserving both spatial attention and spatial working memory ^18–20^. Indeed, LPFC has been shown to contain topographic maps that mirror the organization of visuospatial information in sensory cortex ^21,22^. Although there have been some reports of spatial selectivity in OFC under certain conditions ^23^, most studies have found little OFC spatial coding ^1,10,24,25^. This is broadly consistent with the anatomical connections of these regions. The connections of LPFC are mainly to parietal areas involved in attention, and frontal regions involved in oculomotor and motor control ^26,27^. In contrast, OFC has remarkably few connections to motor regions ^28^, and receives visual input from regions such as the perirhinal cortex ^29^, that are more involved in the representation of objects than the control of spatial attention ^30^. By recording from the same neurons in two different tasks that were specifically designed to differentially tax spatial attention and decision making, we were able to show that the neuronal tuning in OFC and LPFC is remarkably distinct. This does, however, raise the question of how these distinct processes are coordinated.

One possibility is via eye movements. When we first identified the dynamical nature of OFC value representations ^1^ we initially ruled out a relationship with eye movements. However, technical limitations at the time prevented us from recording large numbers of OFC neurons simultaneously. Consequently, our decoding was noisier and often required averaging across trials for its interpretation, potentially obscuring the effects of eye movements in our data. More recent results have shown a strong effect of gaze fixations on value representation. For example, if choice options are presented on a visually cluttered background, they cannot be processed by peripheral vision, a phenomenon known as ‘crowding’ ^31,32^. Instead, they must be fixated sequentially, and under these circumstances OFC represents the value of the option that is fixated ^8^. Our results show that this extends to decision-making, even in the absence of crowding. When subjects were fixating an option, OFC tended to encode the fixated value and LPFC tended to encode the fixated location. Both regions only infrequently represented the unfixated option. When gaze shifted to a new picture, the OFC and LPFC representations shifted accordingly, showing that gaze affected neuronal processing in both regions despite the differences in the kind of information the two areas processed.

Although not as strong as the effect of gaze shifts, we also found a relationship between OFC and LPFC processing as a function of covert attention. Previous groups have demonstrated that the locus of spatial attention can be decoded from population-level neuronal activity in LPFC ^15,16^. In the current study, we used this to reveal an effect of covert processing on OFC value representations over and above that generated by overt gaze shifts, further showing the advantages of using population-level neuronal dynamics to understand cognitive processes that are otherwise difficult to measure in real-time ^33,34^. A critical function of the prefrontal cortex is to allow behavior to be driven by internal representations rather than the sensory environment. Prefrontal damage often results in behavior that is described as ‘stimulus bound’, an inability to avoid reacting or attending to things in this environment ^35–38^. Indeed, if choice options are initially presented but then disappear, subjects will often gaze at the spatial location where the offer appeared. Neurons in OFC represent the value of that offer even though it is no longer present ^39^, supporting the notion that the OFC value code can be related to internal representations.

A key remaining question is how OFC and LPFC processes are coordinated since the two areas are not directly connected ^17^. One possibility is via the ventrolateral prefrontal cortex, since this region has reciprocal connections with both OFC and LPFC ^40–42^, and its neurons show strong tuning both for spatial locations and expected rewards ^43^. In support of this, comparison of neuronal tuning in all three areas during decision-making found that OFC encodes value without respect to the spatial location of offers, and that spatial tuning and the action plan were encoded in ventrolateral prefrontal cortex before the dorsolateral ^25^. Another possibility is that these processes could be coordinated via the thalamus ^44,45^. Neurons in the medial pulvinar are modulated by attentional cues ^46^ as well as parameters relevant to decision-making, such as decision confidence ^47^. Furthermore, medial pulvinar projects to the frontal lobe via two main projections, one to OFC and one to the frontal eye fields ^48^ making it ideal for the coordination of attention and decision-making.

In conclusion, we identified dissociable functions for OFC and LPFC during decision-making encoding value and space, respectively. Despite representing different types of variables, the dynamics related to value and space were correlated. Much of this temporal correlation was explained by gaze since both OFC and LPFC tended to encode the fixated option. However, we found that covert attention could explain additional effects on OFC value dynamics above and beyond the effects of gaze. Thus, while eye movements often reflect the decision-making process, prefrontal neuronal activity also allows decision-making to occur covertly.

## Acknowledgements

This work was funded by NINDS R01-NS116623 and NIMH R01-MH132640 to JDW. The funders had no role in study design, data collection and analysis, decision to publish or preparation of the manuscript.

## Author contributions

NTM and JDW designed the experiment and wrote the manuscript. NTM collected and analyzed the data. JDW supervised the project.

## Methods

### Subjects

All procedures complied with National Research Council guidelines and were approved by the Animal Care and Use Committee at the University of California, Berkeley. Two male rhesus macaques (Subject D and L, respectively) aged 11 and 4 years and weighing 16 and 8.5 kg at the time of recording were used in the current study. Subjects sat in a primate chair (Crist Instruments, Hagerstown, MD), and eye movements were tracked with an EyeLink 1000 system (SR Research, Ottawa, Canada). Stimuli were presented at a viewing distance of 30 cm, and presentation and behavioral conditions were controlled using the NIMH MonkeyLogic toolbox ^49^. Subjects had unilateral recording chambers implanted that allowed access to LPFC and OFC.

### Task design

We trained two monkeys to perform an attention-guided change detection task and a value-based choice task, presented in alternating blocks within each session. Subjects were required to complete a threshold number of trials on the current block (excluding trials that were not successfully initiated or in which the subject failed to make a viable choice response) before advancing to the next block. Based on failure rates from prior sessions, we selected thresholds aiming to yield approximately three blocks each of the attention and value task over a two-hour session, with enough trials per block to adequately balance our task conditions and provide sufficient statistical power to analyze each task condition separately. As such, these thresholds varied between subjects and were occasionally adjusted between sessions, based on each subject’s performance history on each component of the task. Specifically, block criteria were 280 value trials and 230 attention trials for Subject D, and 370 value trials and 240 attention trials for Subject L. On both the attention and value trials, we presented stimuli at 6° eccentricity and at six different locations on the screen (0-300° from the center, at 60° intervals), allowing us to sample across both spatial dimensions. Stimuli in two-item conditions (attention and free choice value trials) always appeared at opposite locations, such that they occupied separate hemifields.

#### Value-based choice task

Value trials were initiated with 500 ms of fixation on a 0.5° central cue. Subjects were then presented with pictures uniquely associated with one of two amounts of juice and one of three reward pay-off probabilities (Figure 1A). On 50% of value trials (free choice), subjects were offered a choice between two pictures associated with different reward prospects; on the other 50% (forced choice), subjects were offered only a single picture. Subjects were allowed to freely view the pictures after stimulus onset, and they selected an option by fixating on it for 350 ms, after which they received a probabilistic reward. All pictures were pre-trained in behavioral training sessions up to 4 days prior to each recording session to encourage stable choice behavior with minimal effects of learning. Subjects were given unique picture sets for each recording.

#### Spatial attention task

Attention trials were initiated with 500 ms of central fixation, after which subjects were presented with one red and one green Gabor grating (Figure 2A). The colors of the gratings cued which target was more likely to rotate after a variable delay period (500-800 ms). Rotations were more frequent at the green or ‘valid’ location (Subject D: 75% of trials, Subject L: 90%) than the red or ‘invalid’ location. Subjects learned to bias attention toward the valid location in anticipation of the change. After subjects held central fixation through the delay, the gratings briefly disappeared for 30 ms and then reappeared, initiating a 500 ms response window. On 75% of trials one of the targets rotated 25°, according to the cue probabilities and subjects were required to saccade to the change location (valid or invalid) to earn a juice reward. The remaining 25% of trials were catch trials, in which neither target rotated. Subjects reported catches by maintaining central fixation through the end of the response window. As we aimed to engage only covert mechanisms of attention, eye movements were prohibited during the delay period. Eye position had to remain within 2° of the fixation cue. Early responses, defined as breaks in fixation before the response period onset or with reaction times below 100 ms, aborted the trial and led to a 500 ms timeout accompanied by a red error screen.

### Behavioral analysis

#### Subjective value models

For the value task, we estimated the subjective values associated with the pictures by fitting decision models to choice behavior on free choice trials. We fit choice behavior for each subject on each session separately, using a softmax decision policy to model the probability that the subject would choose one picture, Option 1, over its alternative, Option 2:

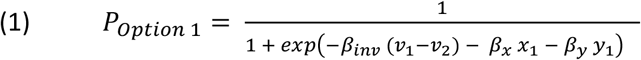

The softmax model includes three free parameters. The inverse temperature, *β*_*inv*_, specifies the stochasticity of choice behavior as a function of the value difference, *v*_*1*_ - *v*_*2*_. The location biases, *β*_*x*_ and *β*_*y*,_ specify the influence of Option 1’s horizontal coordinate, *x*_*1*_, and vertical coordinate, *y*_*1*_, respectively, on choice likelihood.

To estimate value terms *v*_*1*_ and *v*_*2*_, we considered three different subjective value models, which aimed to compute each option’s value by integrating the associated amount, *a*, and probability, *p*, of reward multiplicatively, additively, or both ^50,51^. The expected value (EV) model assumed that subjects were sensitive only to the expected value of each option, as would be expected of an ideal decision-maker optimizing for long-run reward maximization, and was our simplest model with no free parameters:

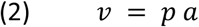

The linear value model treated amount and probability of reward as linearly separable choice variables, allowing us to fit a bias term *β*_*p*_ that captured the subjects’ relative sensitivity to probability over amount during decision-making:

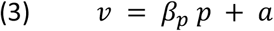

Finally, the hybrid value model incorporated both the multiplicative and additive elements of the other models, with bias terms *β*_*p*_ and *β*_*a*_ to account for sensitivity toward probability and amount, respectively, that cannot be explained by expected value alone:

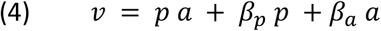

For each session, we used maximum likelihood estimation to fit each of the three subjective value models, along with the softmax decision function, to the subject’s choice behavior. Parameter fits were optimized to maximize the likelihood that the softmax choice probabilities accounted for the subjects’ empirical choices. We performed a formal model comparison using BIC to quantify the relative goodness-of-fit on each session.

#### Signal detection model

We estimated the perceptual sensitivity (*d’*_*val*_ and *d’*_*inv*_) and choice criteria (*c*_*val*_ and *c*_*inv*_) for the valid and invalid targets by fitting change detection behavior to a two-dimensional generalization of the traditional signal detection theory (SDT) model ^52,53^, designed to explain change detection behavior in two-alternative forced choice paradigms such as ours. In this framework, *d’* quantifies how well the decision-maker discriminates the perceptual state associated with a given stimulus change from states associated with no change. The target criterion, *c*, captures bias toward reporting or not reporting a change at each specific target.

The model predicts the probability of each possible response as a function of the stimulus events *X*_*val*_ and *X*_*inv*_, at each respective location (where *X*_*i*_ = 1 for a change at location *i* and *X*_*i*_ = 0 for no change), given the fitted sensitivity and criterion values. The full set of choice probabilities is defined by the following equations, which aim to model the likelihood that a latent decision variable stochastically falls within the decision boundaries, *u*, pertaining to each of the possible choice outcomes within a fixed two-dimensional decision space:

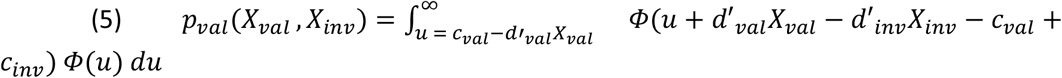

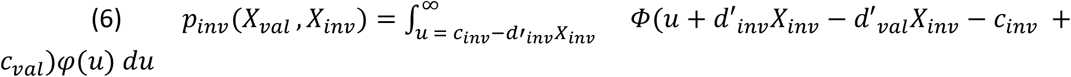

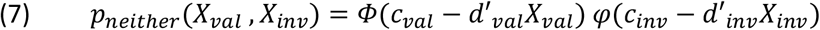

In the formulae above, *φ* and *ϕ* denote the probability density and cumulative distribution functions, respectively, of a Gaussian distribution with unit variance, which we used to model the perceptual signal corresponding to each target.

We fit the model to change detection behavior from each session using maximum likelihood estimation to find the optimal parameter estimates to explain the empirical distributions of response counts (valid, invalid, or neither target) for each of the possible stimulus events (valid change, invalid change, or no change). The fit likelihood was estimated from the trinomial probability distribution of response counts given the model-predicted response probabilities (*p*_*val*_, *p*_*inv*_, and *p*_*neither*_).

#### Eye movement analysis

We defined saccades as eye movements whose speed exceeded 6 standard deviations from the mean velocity during fixation. The raw x- and y-axis velocities were smoothed with a 30 ms Gaussian window prior to saccade detection to correct for noise. The saccade time was defined as the time when the filtered speed peaked after crossing the saccade detection threshold.

We defined fixations during the value-based choice task as periods when the subject’s gaze stayed within a 2° fixation window centered on one of the available pictures. The time of fixation onset was defined as the time when the gaze position first entered this fixation window. Whenever there was a series of consecutive fixations to the same picture, we analyzed only the first of these fixations to avoid double-counting fixations that may have been disrupted due to eye tracking jitter. After quantifying the number of fixations per trial, we categorized each trial as either a single- or multiple-fixation trial. Single-fixation trials were those in which subjects fixated only a single picture without fixating the alternative, whereas multiple-fixation or ‘deliberative’ trials were ones in which subjects switched their gaze from one picture to the other.

### Neurophysiology

Subjects were fitted with titanium head positioners and imaged in a 3T scanner. The resulting images were used to generate three-dimensional reconstructions of each subjects’ skull and brain areas of interest. We then implanted custom radioluminescent recording chambers made of polyether ether ketone (Gray Matter Research). Chambers were placed to allow neurophysiological recordings from OFC and LPFC, targeting the left hemisphere in Subject D and the right hemisphere in Subject L.

For each recording session, we acutely lowered two 45 mm primate Neuropixels probes (IMEC VZW, Leuven, Belgium) into central OFC (areas 11 and 13, between the medial and lateral orbital sulci) and in and around the principal and arcuate sulci of LPFC (areas 8, 9, and 9/46), including both the ventral and dorsal banks of principal sulcus. Electrode trajectories were determined by registering the subject’s MRI into CAD software (OnShape, Cambridge, MA) to design and 3D print custom recording grids which we used to acutely lower the probes (Form 4 3D printer, Form Labs, Somerville, MA). After insertion, the probes were allowed to settle for around 45 - 60 min before the start of the experiment to mitigate drift. We configured the probes to record from 384 active channels in a contiguous block at the tip, allowing dense sampling of neuronal activity along a 3.84 mm span. Neuronal activity was filtered and digitized for action potential bands (300 Hz high-pass filter, 30 kHz sampling frequency) and local field potential (1 kHz low-pass filter, 2.5 kHz sampling frequency). Activity was monitored during experimental sessions and saved to disk using SpikeGLX (https://billkarsh.github.io/SpikeGLX/).

Automatic spike sorting was performed offline using Kilosort4. Because each Neuropixels headstage sampled the data with a slightly different system clock, it was necessary to map all physiological and task-event time series into a common timeline. We accomplished this by broadcasting a 5 Volt, 1 Hz square wave to a unique reference channel on the Neuropixels probes (channel 385) and the non-neural acquisition board. We detected and stored the times of the rising phase of the square wave on each cycle and then used regression to interpolate all events into the timeline of the first Neuropixels probe. To be included for further analysis, neurons had to be present for more than 90% of the experimental session, have a mean firing rate over the course of the entire session >1 Hz, and have fewer than 0.2% of their spikes occurring within 1.2 ms of another neuron’s spikes (that is, <0.2% interspike interval violations).

### Single-neuron tuning

#### Forced choice value trials

For each neuron, we used a series of generalized linear models (GLMs) to quantify the relationship between the firing rate (*FR*) and task variables on each task condition.

On the forced choice trials of the value task, we assessed how neurons encoded the value and location of single pictures through the following GLM:

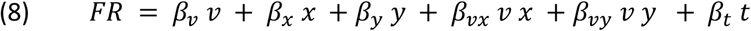

In the above formula, we used the subjective value, *v*, estimated from our behavioral analysis as the value parameter. Spatial parameters *x* and *y* corresponded to the horizontal and vertical coordinates, respectively, where each picture was presented on the screen, and we included nonlinear interaction terms to account for space-value interactions. The timestamp of each trial onset, *t*, was included as a nuisance parameter to account for slow drift in the mean FR.

#### Free choice value trials

We adapted the forced choice GLM to model how neurons encoded picture locations and values on free choice trials, where two competing options were offered:

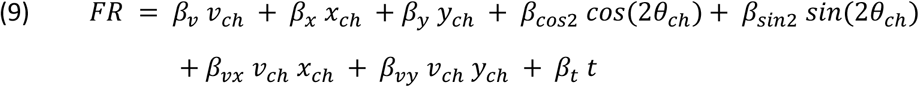

We assessed encoding of value and space pertaining specifically to the chosen picture’s value, *v*_*ch*_, and spatial coordinates, *x*_*ch*_ and *y*_*ch*_, as well as their nonlinear interactions. In addition, we included parameters to model a target-*invariant* spatial response — i.e., a response that varied as a function of the three different axes along which the pictures were presented but did not discriminate between the two possible picture locations on any given axis. We did this by including terms *cos(2θ*_*ch*_) and *sin(2θ*_*ch*_), which define the axis both pictures lie on as a function of the chosen picture’s angular position, *θ*_*ch*_.

#### Attention trials

For each neuron, we fit a GLM to its firing rate that incorporated analogous spatial parameters to those of the free choice GLM:

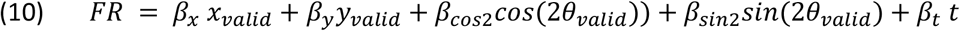

Here the spatial reference frame is defined in terms of the valid target’s location, which is given in Cartesian coordinates *x*_*valid*_ and *y*_*valid*_ and angular position *θ*_*valid*_.

### Population decoding

For each session, we trained linear discriminant analysis (LDA) classifiers to predict picture values (1-6) from either OFC or LPFC population firing rate vectors. Each neuron’s firing rates were first z-scored for each time sample of each task condition. To prevent overfitting due to the high number of neurons relative to training trials, we reduced the dimensionality of the z-scored firing rate data via principal component analysis (PCA). We then selected the first PCs that explained 90% of the variance in the training data and projected both the training and testing data into the PC space identified from the training set.

We first trained a decoder on neural activity during forced choice trials averaged over 100 - 500 ms after picture onset. We then tested the decoder’s performance on free choice activity during the same time window. The decoder was validated using leave-one-out (LOO) cross-validation on all of the forced choice trials to estimate the ‘ground truth’ decoder accuracy.

To investigate single-trial dynamics at a finer temporal resolution, we smoothed firing rates with a 20 ms Gaussian kernel, stepped by 5 ms, prior to z-scoring and PCA. We then trained separate decoders for every time point from 100 - 500 ms after picture onset. We applied these decoders to the corresponding time on free choice trials.

We used LDA classifiers to decode the putative attended location, following a similar procedure. We trained the spatial decoder on z-scored and PCA-transformed firing rate vectors from the delay epoch of the attention task. Decoding was then performed on the firing rates from the value task, which were independently z-scored but projected into the same PC space as the training data. Spatial representations were assessed using the same course-grained fixed windows or fine-grained sliding windows used for value decoding.

### Cross-correlating value and spatial dynamics

To determine whether there was a relationship between the time course of the OFC value representation and the LPFC spatial representation, we cross-correlated the time series of each decoder’s posteriors pertaining to either the fixated or unfixated option for each picture fixation during free choice. We performed the analysis separately for single fixations and follow-up fixations (referred to as ‘switches’) to account for differences in the neural dynamics between the two types of fixations. We excluded sessions where either the chosen value in OFC or chosen location in LPFC were decoded with less than 30% accuracy (D: 6/10 sessions included, L: 4/10 sessions included).

For each decoder on each session and for each subset of fixations, we first subtracted the trial-averaged posteriors to remove any relationships between the average value and spatial dynamics while preserving single-trial interactions. To ensure we had sufficient data to assess correlations, we excluded fixations of < 150 ms duration. Each time series was then binned into 100 ms time bins, which were slid relative to fixation onset in 5 ms intervals. For each fixation, we correlated each bin of value posteriors with each bin of spatial posteriors to yield a trial-wise 2D cross-correlogram between the value and spatial decoders. We then averaged the cross-correlogram across fixations and sessions to quantify the overall relationship between the two decoders. This analysis was restricted to relationships between target-congruent representations (i.e., fixated value versus fixated location or unfixated value versus unfixated location) as the primary question was whether OFC and LPFC encoding was coordinated. In other words, we were testing whether OFC and LPFC represented the same option at the same time and with similar dynamics on a trial-by-trial basis. Significance was assessed using a hierarchical permutation test in which fixation samples for the value and spatial decoders were shuffled relative to each other within each session to derive a null distribution for the mean cross-correlation. We used an alpha level of *p* < 0.01 relative to 5000 null shuffles.

### LFP analysis

LFP signals were analyzed with respect to the following frequency bands: theta (4 - 8 Hz), alpha (8 - 12 Hz), and beta (12 - 30 Hz). We assessed the spectral organization of the LFP signal across each electrode’s recording sites by computing the power spectral density (PSD) on each of the 384 channels (multitaper PSD, N_tapers_ = 4, time-bandwidth product (NW) = 3, sample frequency = 1 kHz). For each frequency band, we selected the channel with the highest average power within each band as the sample channel for the given band on the given electrode, excluding white matter sites (defined based on MRI coordinates and local prevalence of white matter spikes). Theta, alpha, and beta signals were then band-pass filtered from the sampled LFPs, and their phases were extracted using the Hilbert transform.

We asked whether saccades during the value-based choice task were phase-locked to LFP oscillations in any of the frequency bands by regressing the time of the first saccade on each free choice trial against the time of the peak in the first cycle of each band after picture onset. Regressions were performed separately for each subject and each frequency band and were fit using saccade and LFP timing data pooled across sessions.

## Supplemental Figures

**Figure S1.**
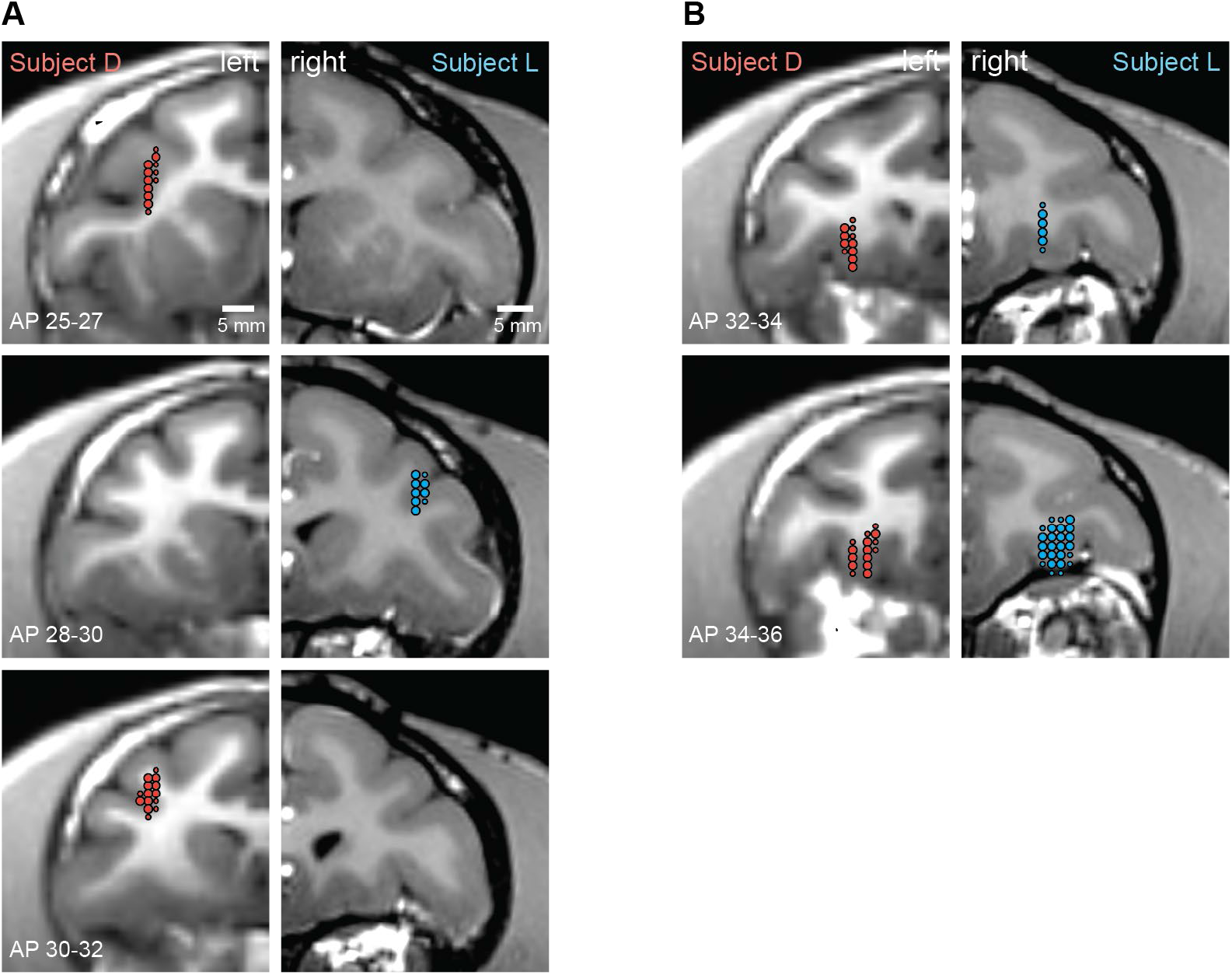
Recording locations. A) LPFC recording locations. Left panels depict Subject D’s recording locations (red) in the left hemisphere; right panels depict Subject L’s recording locations (blue) in the right hemisphere. Small circles denote 5-50 neurons sampled, while large circles denote 50+ neurons. B) OFC recording locations. Same conventions as in (A).

**Figure S2.**
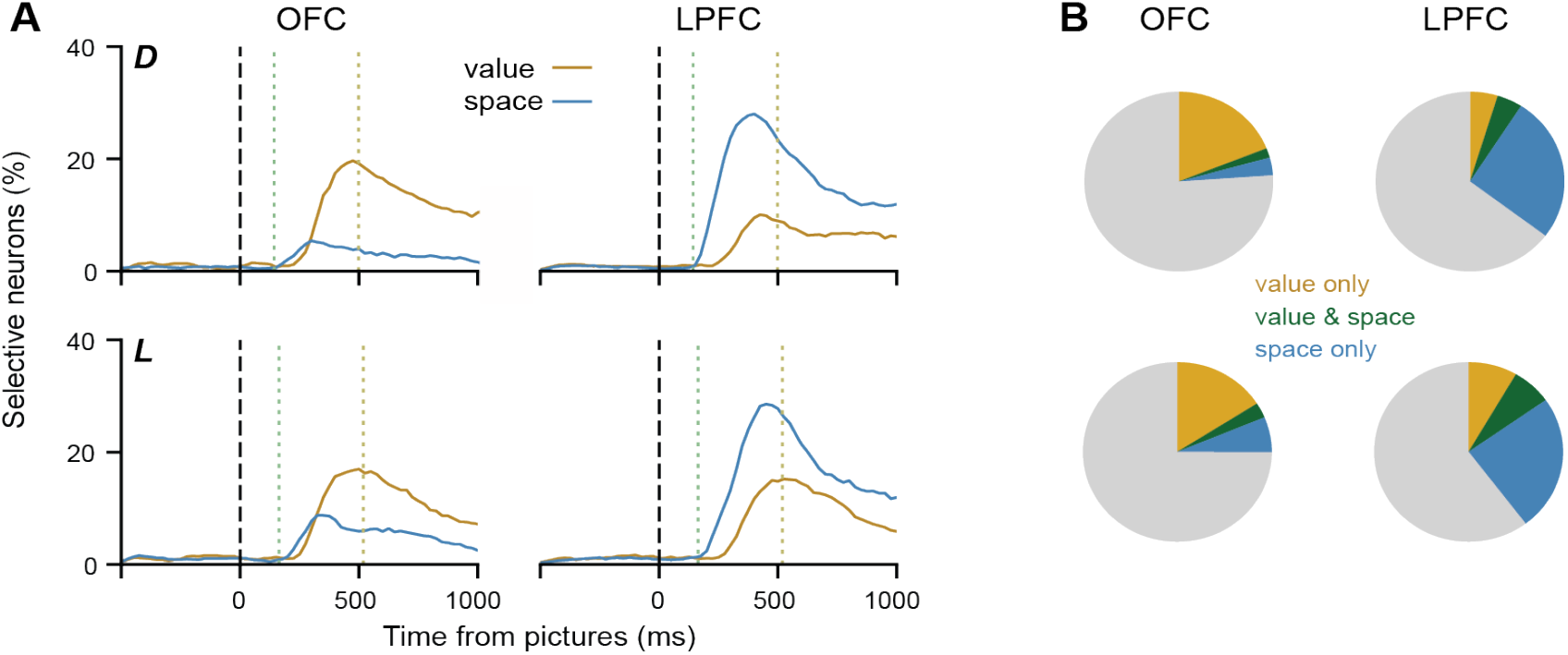
Value and spatial tuning in OFC and LPFC. A) Proportion of single neurons in OFC and LPFC that were identified as selective for the forced choice picture value or spatial location. Selectivity was assessed by fitting a GLM for joint coding of picture values and locations at each time sample (100 ms bins, stepped by 25 ms, *p* < 0.05 for at least 4 consecutive bins). The green dashed line indicates the median saccade reaction time, and the yellow dashed line indicates the median reward acquisition time. B) Proportions of neurons encoding value only, space only, or both as assessed by a fixed-window GLM (100 - 500 ms after picture onset, *p* < 0.01).

**Figure S3.**
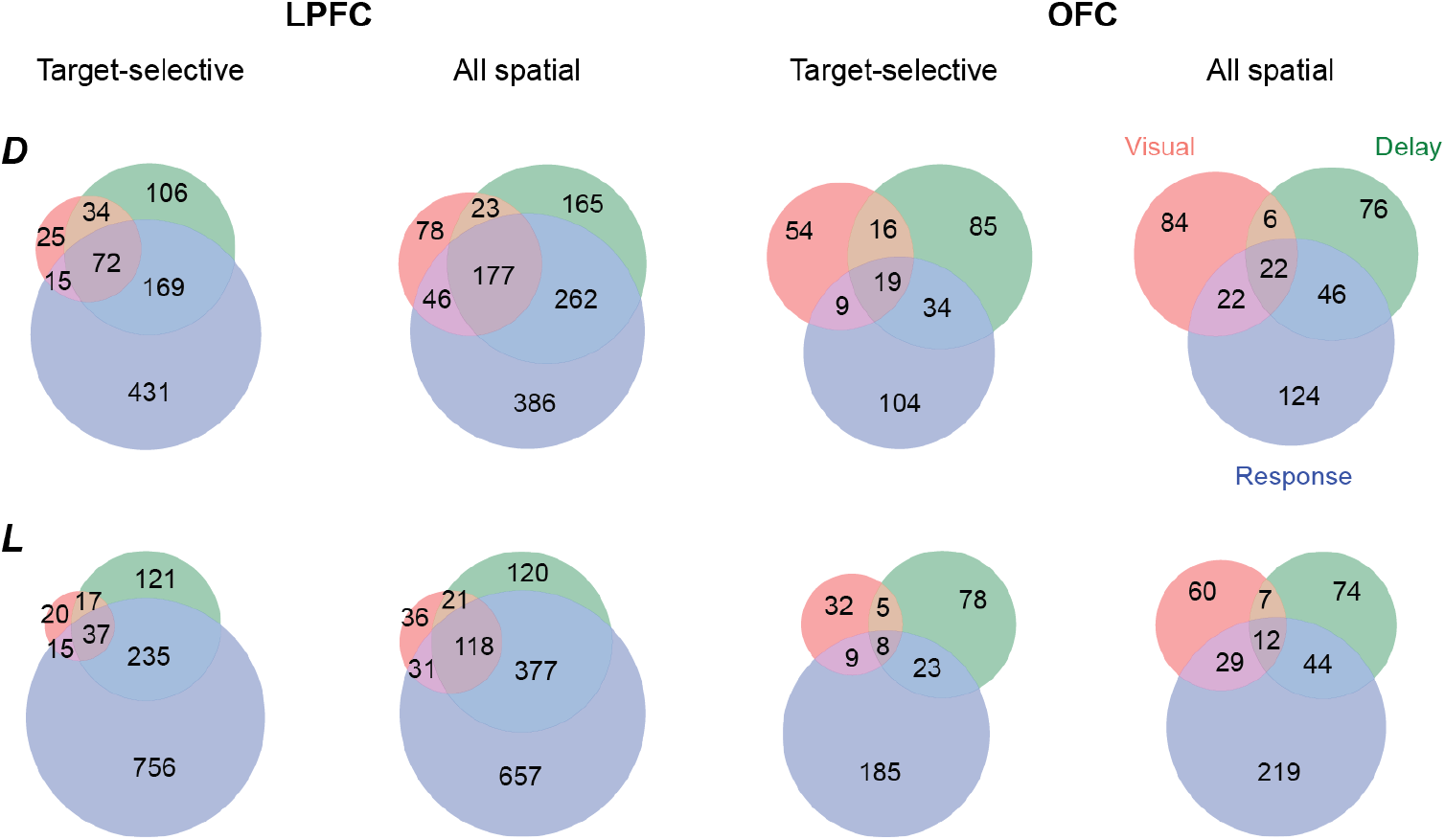
Functional subclasses of attention task spatial neurons. Venn diagrams depicting the number of neurons in LPFC and OFC that significantly encoded space (*p* < 0.01) during the visual, delay, and response epochs of the attention task. Spatial tuning was assessed for target-selective spatial coding (encoding the target location specifically) as well as for all spatial coding (counting either target-selective or target-invariant coding as spatial tuning).

**Figure S4.**
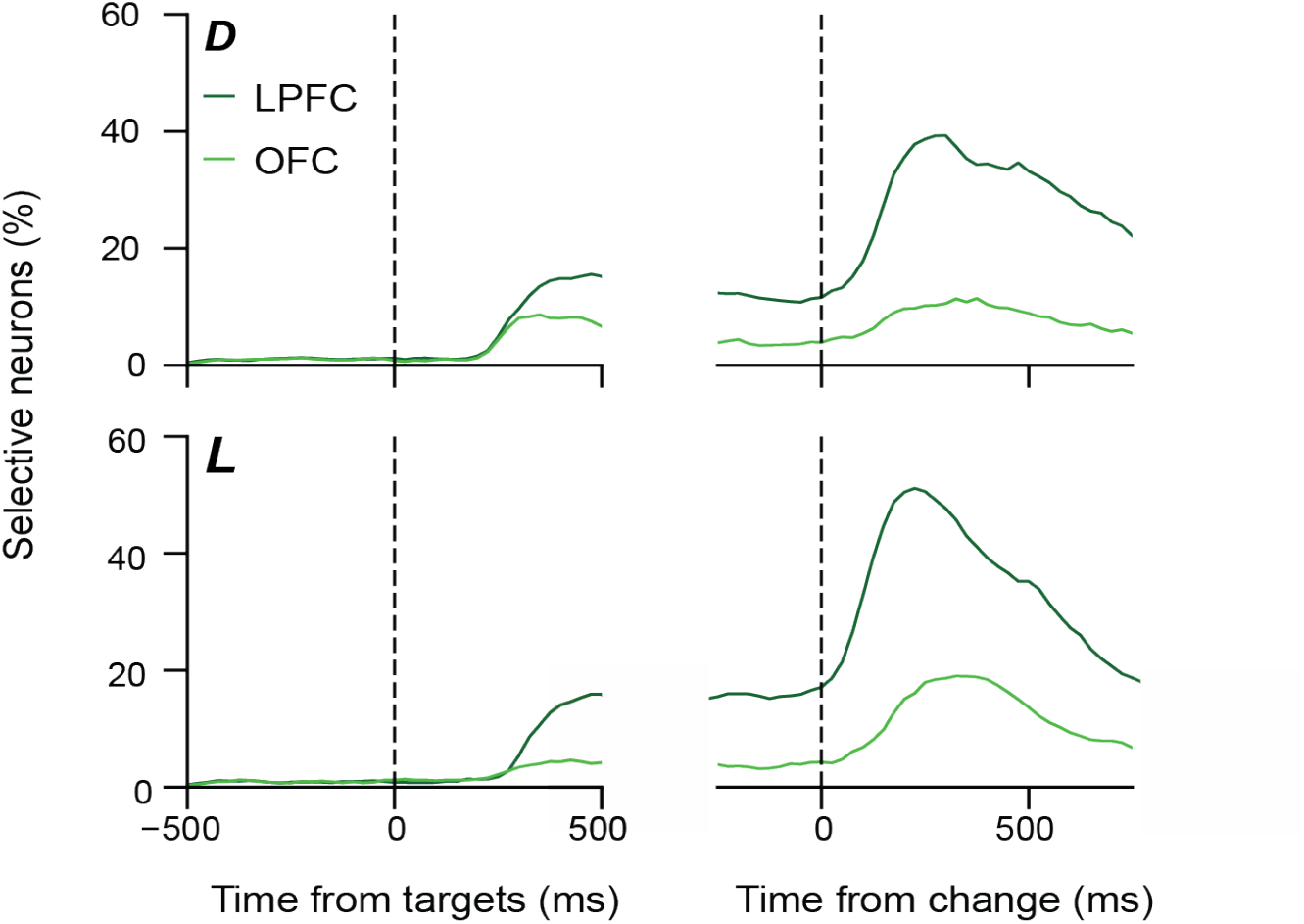
Spatial tuning during the attention task. Proportion of single neurons in LPFC and OFC encoding target location, assessed by fitting a GLM for spatial coding at each time sample (100 ms bins, stepped by 25 ms, *p* < 0.05 for at least 4 consecutive bins).

